# Evaluation of machine learning methods to predict peptide binding to MHC Class I proteins

**DOI:** 10.1101/154757

**Authors:** Rohit Bhattacharya, Ashok Sivakumar, Collin Tokheim, Violeta Beleva Guthrie, Valsamo Anagnostou, Victor E. Velculescu, Rachel Karchin

## Abstract

Binding of peptides to Major Histocompatibility Complex (MHC) proteins is a critical step in immune response. Peptides bound to MHCs are recognized by CD8+ (MHC Class I) and CD4+ (MHC Class II) T-cells. Successful prediction of which peptides will bind to specific MHC alleles would benefit many cancer immunotherapy appications. Currently, supervised machine learning is the leading computational approach to predict peptide-MHC binding, and a number of methods, trained using results of binding assays, have been published. Many clinical researchers are dissatisfied with the sensitivity and specificity of currently available methods and the limited number of alleles for which they can be applied. We evaluated several recent methods to predict peptide-MHC Class I binding affinities and a new method of our own design (MHCnuggets). We used a high-quality benchmark set of 51 alleles, which has been applied previously. The neural network methods NetMHC, NetMHCpan, MHCflurry, and MHCnuggets achieved similar best-in-class prediction performance in our testing, and of these methods MHCnuggets was significantly faster. MHCnuggets is a gated recurrent neural network, and the only method to our knowledge which can handle peptides of any length, without artificial lengthening and shortening. Seventeen alleles were problematic for all tested methods. Prediction difficulties could be explained by deficiencies in the training and testing examples in the benchmark, suggesting that biological differences in allele-specific binding properties are not as important as previously claimed. Advances in accuracy and speed of computational methods to predict peptide-MHC affinity are urgently needed. These methods will be at the core of pipelines to identify patients who will benefit from immunotherapy, based on tumor-derived somatic mutations. Machine learning methods, such as MHCnuggets, which efficiently handle peptides of any length will be increasingly important for the challenges of predicting immunogenic response for MHC Class II alleles.

**Author Summary:** Machine learning methods are a popular approach for predicting whether a peptide will bind to Major Histocompatibility Complex (MHC) proteins, a critical step in activation of cytotoxic T-cells. The input to these methods is a peptide sequence and an MHC allele of interest, and the output is the predicted binding affinity. MHC Class I and II proteins bind peptides of 8-11 amino acids and 16-26 amino acids respectively. This has been an obstacle for machine learning, because the methods used to date can only handle fixed-length inputs. We show that a recently developed technique known as gated recurrent neural networks can handle peptides of variable length and predict peptide-MHC binding as well or better than existing methods, at substantially faster speeds. Our results have implications for the hundreds of MHC alleles that cannot be predicted with current methods.

## Introduction

The presentation of peptides bound to major histocompatbiity complex (MHC) proteins on the surface of antigen-presenting cells and subsequent recognition by T-cell receptors is fundamental to the mammalian adaptive immune system. Recent advances in cancer immunotherapy have highlighted the need for improved understanding of which peptides will bind to MHC proteins and generate an immune response [1], [2], [3], [4]. In particular, peptides that harbor somatic mutations specific to a patient’s tumor, known as neoantigens, can inform treatment [5] [6]. Because experimental characterization of peptide-MHC binding is costly and time-consuming, computational researchers have been working for decades on *in silico* tools to predict peptide-MHC affinities [7]. Approaches have included sequence-based profiles [8] [9], structure-based predictions [10], generative probabilistic models [11], and machine learning [12] [13] [14].

Neural network supervised machine learning methods are the most widely-used technique and have been shown to outperform other methods in multiple studies [15] [16] [17]. However, many researchers remain dissatisfied with the available *in silico* peptide-MHC binding predictors for purposes of neoantigen discovery, particularly in a clinical setting [18] [19].

Neural networks were originally modeled on the human brain. They consist of connected units, organized into layers. Theoretically, given enough layers and units, they can act as universal function approximators [20]. However, as the number of units increases, they become prone to overfitting (reviewed in [21]). Most recently, deep learning research has introduced innovative network architectures and regularization techniques, allowing for training of very deep and wide networks to approximate complex functions, with reduced risk of overfitting. Given sufficient training data, deep architectures outperform other methods in many domains [22]. They are now widely used in robotics, self-driving vehicles, sentiment classification, machine translation, and user-based recommendation systems [23] [24] [25] [26] [27].

In this work, we rigorously assessed the performance of the most widely-used and several recently published machine learning methods to predict peptide-MHC binding: NetMHC [28], NetMHCpan [29], MHCflurry [30], SMMPMBEC [15], HLA-CNN [13], and a new method designed by us called MHCnuggets. The methods include standard neural networks, a Bayesian matrix method, and deep learning methods. Evaluation was done on a carefully designed and previously published benchmark set of immuno-fluorescent binding experiments for diverse peptide-MHC I allele pairs [16], derived from the IEDB database [31]. The benchmark set was cleaned for redundancy, using a protocol designed by the MHCflurry team.

Our results indicate that four methods: NetMHC, NetMHCpan, MHCflurry, and our new method MHCnuggets have best-in-class prediction performance, and of these methods MHCnuggets was substantially faster. MHCnuggets is a gated recurrent neural network, which can handle peptides of any length. This is a methodologically important advance, because current machine learning methods must either artificially shorten or lengthen peptides, or train a separate classifier for each peptide length. These strategies will become intractable as the field expands to the important problem of handling MHC Class II alleles [32], which bind peptides of highly variable lengths.

MHC proteins are highly polymorphic, and there are thousands of MHC Class I alleles, each with a different peptide binding surface. While there are tens of thousands of experimentally characterized binding affinities in the benchmark set, they are distributed unevenly across the alleles. For the most common alleles, particularly those associated with European ancestry, there are as many as 9000 characterized peptides, while for others there may be as few as 100. When we stratified prediction performance by individual alleles, performance was seen to vary widely by allele. Most of the variance in prediction performance could be explained by data-driven differences between alleles in the benchmark set: the number of training examples, imbalance between binders and non-binders in training and test sets, and sequence diversity of the experimentally characterized peptides in the training set. This result contradicts the previously published hypothesis that differences in prediction performance are primarily due to biochemical differences between allele-specific peptide binding properties [33].

## Results/Discussion

### Prediction of peptide-MHC Class I binding affinities

We compared five previously published prediction methods and a new method of our own design. Predictors were rigorously assessed by measuring their performance on experimental IC50 measurements for peptide-MHC pairs, covering 51 alleles [16]. We selected the 51-allele benchmark because it is more informative than one based on fewer MHC alleles. Machine learning methods must be evaluated by their ability to make correct predictions on new data, which requires that examples are separated into non-overlapping training and test sets. The predictors were trained on a set of peptide-MHC pairs with no overlap to the test set. To the best of our ability, the predictors were trained on the same peptide-MHC pairs, but NetMHC and NetMHCpan do not make their training software available, so we used their self-reported results. These methods have been previously reported to augment their training sets with additional proprietary data, beyond what is in the benchmark training set used in this work [16, 30], so their self-reported performance estimates may be overly optimistic.

**Table 1.** Prediction performance of the six tested methods. Three metrics were applied. Continuous-valued predictions of peptide-MHC binding affinity were assessed wih Kendall-Tau cor-relation (K-Tau). Classifications of peptides as binders or non-binders (at IC50 threshold of 500nM) were assessed with F1 score and AUC (Methods and Materials). The best-peforming method according to each metric is highlighted in bold.

|  | **K-Tau** | **F1** | **AUC** |
| --- | --- | --- | --- |
| MHCnuggets | **0.589** | **0.810** | 0.931 |
| NetMHC | 0.588 | 0.808 | 0.932 |
| MHCflurry | 0.587 | 0.785 | **0.933** |
| NetMHCpan | 0.584 | 0.803 | **0.933** |
| SMMPMBEC | 0.578 | 0.791 | 0.921 |
| HLA-CNN | 0.480 | 0.750 | 0.874 |

We found that MHCnuggets, NetMHC, MHCflurry, and NetMHCpan had the best overall prediction performance (Table 1). Three metrics were applied. Continuous-valued predictions were assessed with the Kendall-Tau coefficient, which measures correlation of predicted and experimental IC50 values. The continuous values were thresholded to produce a classification for each peptide (binder or non-binder). We used an IC50 threshold of 500nM (binder ≤ 500nM), for which there is strong biological support [34]. The classifications were assesed with the F1 score and area under the ROC curve (AUC). The F1 score measured precision and recall at the IC50 500nM threshold, giving equal weight to each. The AUC metric was more forgiving, because it considered all possible thresholds, and a predictor can have good AUC based on its classification at biologically uninformative thresholds. By considering all three metrics, we produced a more comprehensive assessment than by using only one. However, Kendall-Tau and F1 scores provide more stringent assessments of prediction performance than AUC.

To further compare the predictions of the tested methods, we produced a list of peptide-MHC pairs, ranked by each method’s predicted binding affinities. The ranked predictions were strikingly correlated (Spearman rank correlation ≥ 0.9 for all method pairings, with the exception of HLA-CNN) (Figure 1). Given that the methods used different training algorithms and network architectures, the high rank correlation on the level of individual predictions was surprising. This result means that ranking of peptide-MHC affinities is similar among the best methods identified by us. If predictors are used only to rank peptides, any of these methods will produce equivalent results. However, the methods’ classification of peptides as binders or non-binders will not necessarily yield equivalent results.

**Figure 1.**
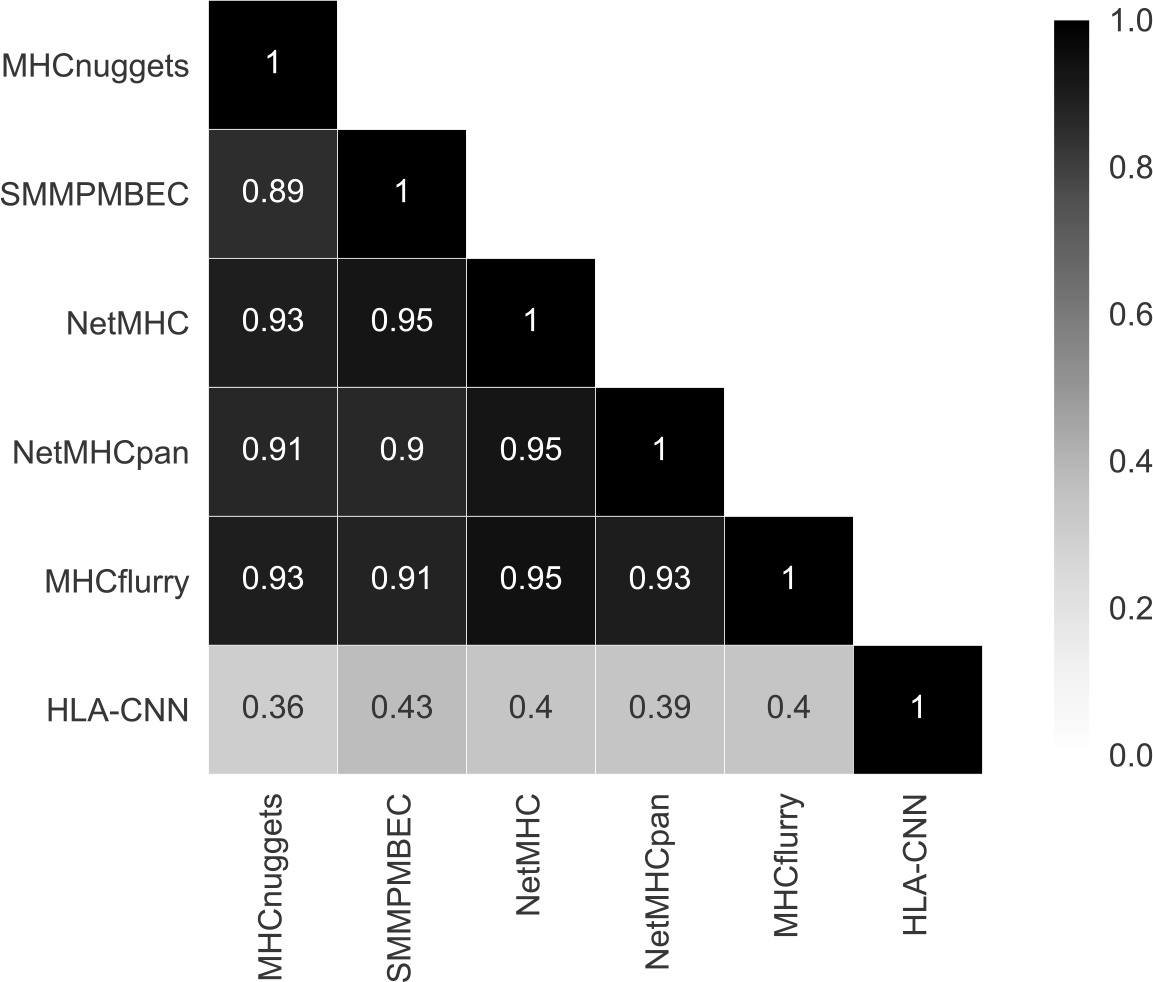
Spearman rank correlation of predicted peptide-MHC binding affinities by six methods.

Convolutional neural networks (CNNs), a popular deep learning technique did not do well in our assessment. HLA-CNN had the lowest predictive performance and the lowest Spearman rank correlation with other methods. In addition, CNNs developed by our group did not perform as well as other neural network architectures (Supplementary Information). Peptide-MHC binding is highly position-sensitive [35], while CNNs were designed to handle tasks where position invariance is important. For example, they excel at object detection, where the same object might be present at different locations in an image. In contrast, a valine amino acid residue at position six of a peptide may have very different impact on binding than one at postion nine. Our results suggest that CNNs are not well suited to the problem of peptide-MHC binding.

Finally, we assessed the speed of each method at the same prediction task. We computed the runtime for each method to predict the binding affinities of all peptide-MHC pairs in the test set (163,898), with respect to a single MHC allele (HLA-A*02:01). Each run was repeated five times and averaged. Runtimes varied substantially (Table 2). SMMPMBEC had the best runtime, followed by HLA-CNN. MHCnuggets was substantially faster than MHCflurry, NetMHC, and NetMHCpan, and NetMHCpan was the slowest method by a large margin. Timing was done on a single compute node containing 2 Intel Xeon E5-2680v3 (Haswell) processors running at 2.5GHz, with 12 cores and 128 GB RAM.

**Table 2.** Runtime analysis. Average runtime for each predictor on peptides in the test set and estimated of the runtime on the Cancer Genome Atlas head and neck cancer samples.

| Method | Test set runtime [sec] | Estimated TCGA HNSC runtime [min] |
| --- | --- | --- |
| SMMPMBEC | 1.25±0.14 | 1.56 |
| HLA-CNN | 9.82±0.083 | 12.24 |
| MHCnuggets | 22.76±0.50 | 23.38 |
| MHCflurry | 49.12±1.33 | 61.24 |
| NetMHC | 80.89±10.3 | 100.85 |
| NetMHCpan | 282.58±18.64 | 352.29 |

We estimated how these runtime differences would scale to an application in which a cohort of cancer patients was evaluated for predicted neoantigens, based on somatic mutations called from whole-exome sequencing. As an example, we considered, the head and neck cancer (HNSC) samples in the Cancer Genome Atlas [36] (495 samples as of June 15, 2017). For each sample, there are on average 4128 candidate peptides (considering all possible 8-11 length windows around each somatic missense mutation) and six potential MHC Class I alleles (3 loci, possibly all heterozygous).

We did not include two recent methods (sNebula [37], PSSMHCpan [38]), because they have not made their training software available or published self-reported results on the benchmark test set.

### Allele-specific differences in *in silico* prediction performance are largely data-driven

Prediction performance varied substantially by MHC allele. Kendall-Tau correlations for the HLA-A*25:01, HLA-A*80:01, HLA-B*18:01, and HLA-B*45:01 alleles were particularly low, while those for H2-Db, HLA-A*02:03, HLA-B*15:01, HLA-A*02:02, HLA-A*29:02, HLA-A*02:01, HLA-A*68:02, HLA-A*03:01, and HLA-A*11:01 were high (Figure 2, Data S1). There were a few examples where one of the methods had superior performance for a particular allele (NetMHCpan F1 score for HLA-A*25:01, MHCnuggets F1 score for HLA-B*08:02). More strikingly, in Figure 2, the alleles could be divided into two groups, 34 for which all the methods performed well (except for HLA-CNN) (top two-thirds of the rows) and 17 for which none performed well (bottom one-third of the rows).

**Figure 2.**
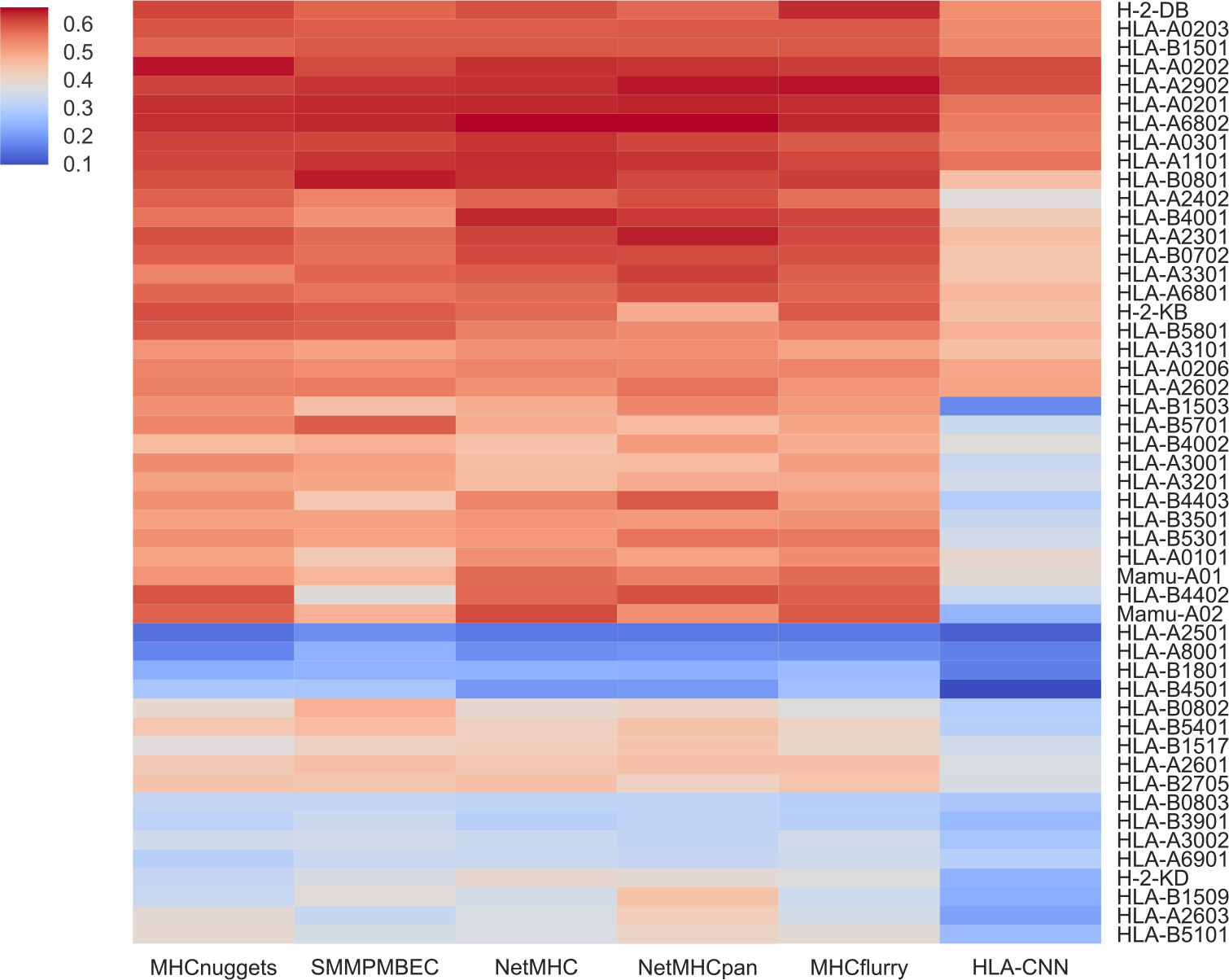
Prediction performance for 51 MHC Class I alleles. Hierarchically clustered heat map of methods and alleles shows performance by Kendall-Tau correlation (high=red, low=blue). The alleles separate into a group for which all methods except HLA-CNN perform well (top two-thirds of the rows) and those for which none peform well (bottom one-third of the rows). Raw Kendall-Tau correlations are in Data S1.

We considered whether properties of the data used to train and test the methods could explain the variation in prediction performance for different MHC Class I alleles. ANOVA comparison of nested linear regression models was used to assess the variance in prediction performance. Given the lack of correlation between HLA-CNN and the other methods coupled with its weak prediction performance, it was not included in the analysis. Using Kendall-Tau correlation or F1 score as the response variable, we identified several significant covariates (p<0.05). These were the number of training examples (peptides with measured binding affinities) for each allele; imbalance between binder and non-binder examples in the training or test sets; and peptide sequence diversity in the training examples (measured by network modularity). Details are in Materials and Methods.

Our results suggest that properties of the training and test data explain a large proportion of the variance among performance of methods across different MHC Class I alleles. For performance measured by Kendall-Tau correlation, 69.2% of the variation across alleles was explained by test examples imbalance (65.4%, p=4.8e-59) and number of training examples (+3.8%, p=6.8e-08). For performance measured by F1 score, 60.9% of the performance variation was explained by training example imbalance (40.1%, p=2.1e-29), number of training examples (+9.3%, p=1.17e-10), binder peptide sequence diversity in the training examples (+9.3%, p=1.6e-06), sequence diversity of all peptides in the training examples (+4.5%, p=4.53e-05), and test examples imbalance (+4%, p=1.03e-6). Choice of prediction method did not explain any additional variance in performance.

It has been previously suggested that allele-specific biochemical properties of peptide-MHC complexes were responsible for alleles whose peptide affinities are not well predicted [33]. Given the ANOVA results presented here, it is more likely that differences in prediction performance are data-driven. We consider this to be an optimistic conclusion, because it suggests that as larger and more diverse sets of peptides are experimentally characterized, with respect to more MHC alleles, prediction methods will substantially improve.

## Materials and methods

### Data collection

The Immune Epitope Database (IEDB) [31] provides a large public set of experimentally characterized peptides and peptide-MHC binding affinities. Database entries were curated from published literature, and the majority of affinities were calculated based on immunofluorescent assays. These affinities are represented as an IC50 value, the half-maximal inhibitory concentration in nano-molar (nM) units of peptide to MHC molecules. A total of approximately 250,000 examples from immunofluorescent assays was available as of May 2017, spanning multiple mammalian and avian species, and 740 MHC alleles, of which 459 are MHC Class I.

For purposes of benchmarking *in silico* peptide-MHC Class I binding predictors, Kim et al. generated a new dataset from IEDB, partitioned into training and test sets [16]. The dataset was further processed by a shell script [30] and available at https://github.com/hammerlab/mhcflurry/tree/master/downloads-generation/data_kim2014, that removed any peptide in the test set with identical length and ≥ 80% sequence identity to a peptide in the training set. The resulting benchmark contains 106 unique MHC alleles and 137,654 IC50 measurements, published prior to 2009 (training set) and 51 unique MHC alleles with 26,888 IC50 measurements, published from 2009-2013 (test set). Two alleles (HLA-B*46:01, HLA-B*27:03) did not contain any peptides defined as binders in this work (IC50<500nM) and were dropped from the analysis. All peptides in the benchmark set consist of 8-11 amino acid residues.

### MHCnuggets

MHCnuggets uses a gated recurrent unit neural network architecture (GRU) [39] and is trained on IC50 values from immuno-fluorescent binding experiments for peptide-MHC Class I pairs. Peptides are represented to the network as a series of amino acids; each amino acid is represented as a 21-dimensional smoothed, one-hot encoded vector (0.9 and 0.005 replace 1 and 0). The GRU accepts inputs of any length, so no cutting or padding of peptides is needed.

A separate network was trained for each MHC allele, and a transfer learning protocol (Figure S1) was applied. First, we trained a network for the allele with the largest number of training examples (HLA-A*02:01). This trained network contained approximately 21,000 weight parameters. The weights were used to initialize and train the networks for all other MHC alleles. Next, we assessed the prediction performance of each network on the training examples for each of the alleles. For each allele, if the network that performed best was not the HLA-A*02:01 network, we did a second round of training, with the best performing network’s weights used in the intialization step (Figure S2). The prediction performance for alleles in the benchmark test set consistently improved when transfer learning was applied (Figure S3).

All networks were implemented with the Keras python package (TensorFlow back-end) [40] [41]. Each network has a fully connected layer of 64 hidden units and a final output layer of a single sigmoid unit, related to the predicted binding affinity as *y* = *max*(0, 1 *− log*_50*K*_ IC50). Networks were trained for 200 epochs, using backpropagation with the Adam optimizer [42], and a learning rate of 0.001. Regularization was performed with dropout and recurrent dropout [43] probabilities of 0.2. The number of hidden units, dropout rate [44], and number of training epochs was estimated by three-fold cross-validation on the HLA allele with the largest number of entries (HLA-A*02:01).

MHCnuggets software is available at https://github.com/KarchinLab/mhcnuggets.

### Comparison metrics

#### Kendall-Tau correlation

The Kendall-Tau correlation is defined as

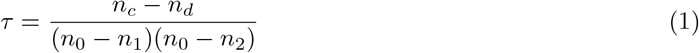

where *n_c_* is the number of concordant pairs, *n_d_* is the number of discordant pairs, and *n*_0_, *n*_1_, and *n*_2_ are given by Equations 2, 3, 4.

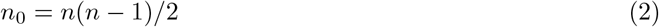

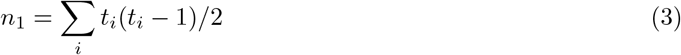

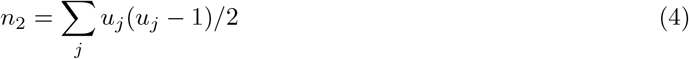

where *n* is the total number of experimental and predicted binding affinity pairs, *t_i_* is the number of tied values in the *i^th^* group of tied experimental affinities, and *u_j_* is the number of tied values in the *j^th^* group of tied predicted affinities.

#### F1 score

The F1 score is defined as

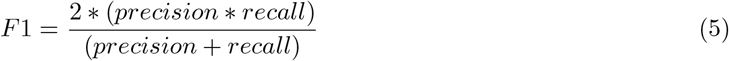

where 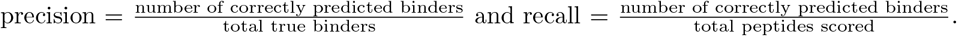

#### Area under the receiver operating characteristic curve

The true positive rate (precision) and false positive rate 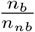 is computed at each possible score threshold, and the final metric is the area under the resulting curve.

### Allele-specific performance covariates

For each of the 51 MHC Class I alleles in the benchmark test set, we calculated the following covariates: the number of training examples *n*, the imbalance between binders and non-binders in the training examples 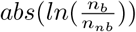 where *n_b_* is the number of binder training examples ≤ 500nM and *n_nb_* is the number non-binders > 500nM (in practice we use *abs* 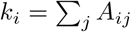 to handle cases where *n_b_ > n_nb_*), the sequence diversity of the training example peptides (calculated as network modularity as described below), the number of test examples, and the imbalance between binders and non-binders in the test examples. The ANOVA R package was utilized to measure the variance in prediction performance of the tested methods explained by each covariate, using nested linear regression models and with either F1 score or Kendall-Tau correlation as the response variable. Covariates were added to each model with a greedy maximization of variance algorithm, in which the covariate that increased the variance the most at each iteration was added to the model, if it had a p-value < 0.05. HLA-CNN was not included in the analysis (as described in Results).

### Network analysis of peptide diversity

We constructed a network for each of the 51 MHC Class I alleles in the benchmark test set. The nodes of these networks were the peptide sequences found in the training set of each allele. Two nodes were connected by an edge if the sequence identity between the peptides was > 0 (*i.e*., they shared at least one identical amino acid residue at the same position). Edges were weighted by sequence identity (normalized by peptide length). Peptides were trivially aligned by cutting and padding them to length nine, as described in (MHCnuggets). Community detection was performed with the fast unfolding algorithm [45], as implemented by the `community_multilevel` function in *igraph*. Once nodes were assigned to communities, the weighted modularity of the community assignment was calculated as:

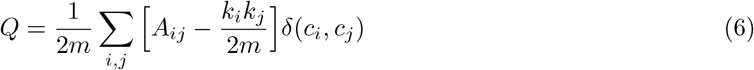

where *A_ij_* is the weight of the edge between *i* and *j*, *k_i_* =Σ_*j*_A_ij_is the sum of the weights of the edges associated with vertex *i*, *c_i_* is the community to which vertex *i* is assigned, δ is an indicator function such that δ(*u, v*) = 1 if *u* = *v* and is 0 otherwise, and 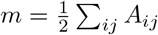

## Acknowledgments

We would like to thank the MHCflurry team for their inspiration and the obvious influence on the name of our methods, MHCnuggets. MHCflurry open sources all of their software and data, which makes benchmarking studies such as this one possible. The results shown here are in part based upon data generated by the TCGA Research Network: http://cancergenome.nih.gov.

## Supplementary Information

### Tested methods to predict peptide-MHC Class I binding affinities

#### NetMHC/NetMHCpan

NetMHC [28] is a single-layer fully connected neural network that encodes nine amino acid residue peptides with a 378-length input vector, which incorporates both a smoothed one-hot encoding (0.9 and 0.05 replace 1 and 0) and a BLOSUM-62 encoding of each amino acid. To accommodate peptides that are shorter or longer than nine residues, contiguous padding or cutting operations are applied at every possible position. When padded or cut versions of the same peptide receive different predicted binding affinities, the strongest affinity is selected. Separate networks are trained for each MHC allele.

NetMHCpan [29] is also a single-layer fully connected neural network that encodes both peptide and polymorphic residues of the relevant MHC allele. It uses the same smoothed one-hot plus BLOSUM-62 encoding and padding/cutting protocol as NetMHC. A single network is trained for all MHC alleles. NetMHC and NetMHCpan are currently the most widely used in silico peptide-MHC I binding tools.

Both methods use artificially generated non-binding peptides, by applying the NetChop algorithm [46] to the entire human proteome.

#### MHCflurry

MHCflurry [30] is a neural network, which jointly discovers informative amino-acid residue encodings and predicts peptide-MHC I binding affinities. Each amino acid residue type is assigned an integer, and peptides are encoded as 9-length vectors of integers. An initial embedding layer maps input peptides to a 32-dimensional space, which then feeds into a fully connected layer. Padding and cutting of peptides that are shorter or longer than 9 residues is done identically to the protocol of NetMHC/NetMHCpan. The final predicted binding affinity is the geometric mean of all padded and cut versions of the same peptide. MHCflurry augments its training data with peptides randomly generated *in silico* from a uniform distribution.

#### SMMPMBEC

SMMPMBEC [15] is a Bayesian framework for regularized least-squares regression. Peptides to be used for training are represented with one-hot encoding and stacked to form a single NxL matrix, where N is the number of peptides and L is the peptides’ length x 20. A scoring matrix is trained to minimize the error between N experimental binding affinities and the matrix product of the peptide matrix and the scoring matrix. The optimization is constrained by a pre-trained 20x20 pairwise substitution matrix for the amino acids, based on the covariance of their contributions to peptide-MHC I binding free energy in different contexts. A scoring matrix is computed for each MHC allele and each peptide length. To predict the affinity for a peptide of interest, its one-hot representation is multiplied by the scoring matrix.

#### HLA-CNN

HLA-CNN [13] is a deep learning convolutional neural network, consisting of an embedding layer, two 1D convolutional layers, and a fully connected layer. The input to the embedding layer is an Lx15 matrix, which is initialized with a learned peptide representation based on the natural language processing (NLP) skip-gram model [47]. Skip-gram is an unsupervised technique to discover word embeddings. Sentences are encoded as vectors of integers and projected into a lower dimensional space. HLA-CNN treats peptides as sentences and amino acid residues as words. The initial embedding is learned from the set of all peptides across all MHC alleles in the training set. The convolutional layers consist of 32 filters, with kernel size of 7, stride=1, and are initialized with Glorot normal distribution [48]. A network is trained for each MHC allele and each peptide length.

#### Additional neural network methods developed for this paper

While developing MHCnuggets, we experimented with long short-term memory network (LSTM) [49], convolutional neural network (CNN) [50], and standard neural network architectures. The LSTM had slightly worse prediction performance than the GRU architecture (Kendall-Tau 0.587, F1 score 0.806, AUC 0.931) and was slightly slower (test set runtime 23.91 sec ±0.37). It also accepts variable length inputs. We consider the LSTM to be a viable alternative for future work. The CNNs had the worst prediction performance of the tested methods. This network architecture is designed to fit weights using multiple sliding windows (kernels) across each peptide. First, we tried kernels that covered two or three amino acid residues (Kendall-Tau 0.447, F1 score 0.64, AUC 0.845). We also tried adding a kernel that covered the full length of the peptide, which improved performance (Kendall-Tau 0.563, F1 score 0.795, AUC 0.918), but did not match the performance of the GRU or LSTM. CNNs require a fixed length input. Based on our observation that peptide cutting and shortening protocols substantially slowed runtime, we applied a simplified protocol in which any peptides that were longer or shorter than 9 residues were cut or padded once. To ensure that padding and cutting were sensitive to primary and secondary anchor residue positions, cutting and padding were restricted to positions 6 or 7. With this protocol, the CNNs were among the fastest methods tested (9.78 ±0.116 sec for the two kernel CNN and 7.13 ±0.073 sec for the three kernel CNN on the benchmark test set). Finally, we tried a network similar to NetMHC, a fully-connected single-layer network that required fixed-length inputs, and applied the same simplified cutting and padding procedure used by the CNNs. The standard neural network performed well, with the best F1 score of any method but a lower Kendall-Tau than the GRU and LSTM architectures (Kendall-Tau 0.581, F1 score 0.814, AUC 0.931). It was also among the fastest methods (7.32 ±0.053 sec on the test set). However, in contrast to the recurrent neural networks, it does not offer any methodological advances over the other tested methods.

**Figure S1.**
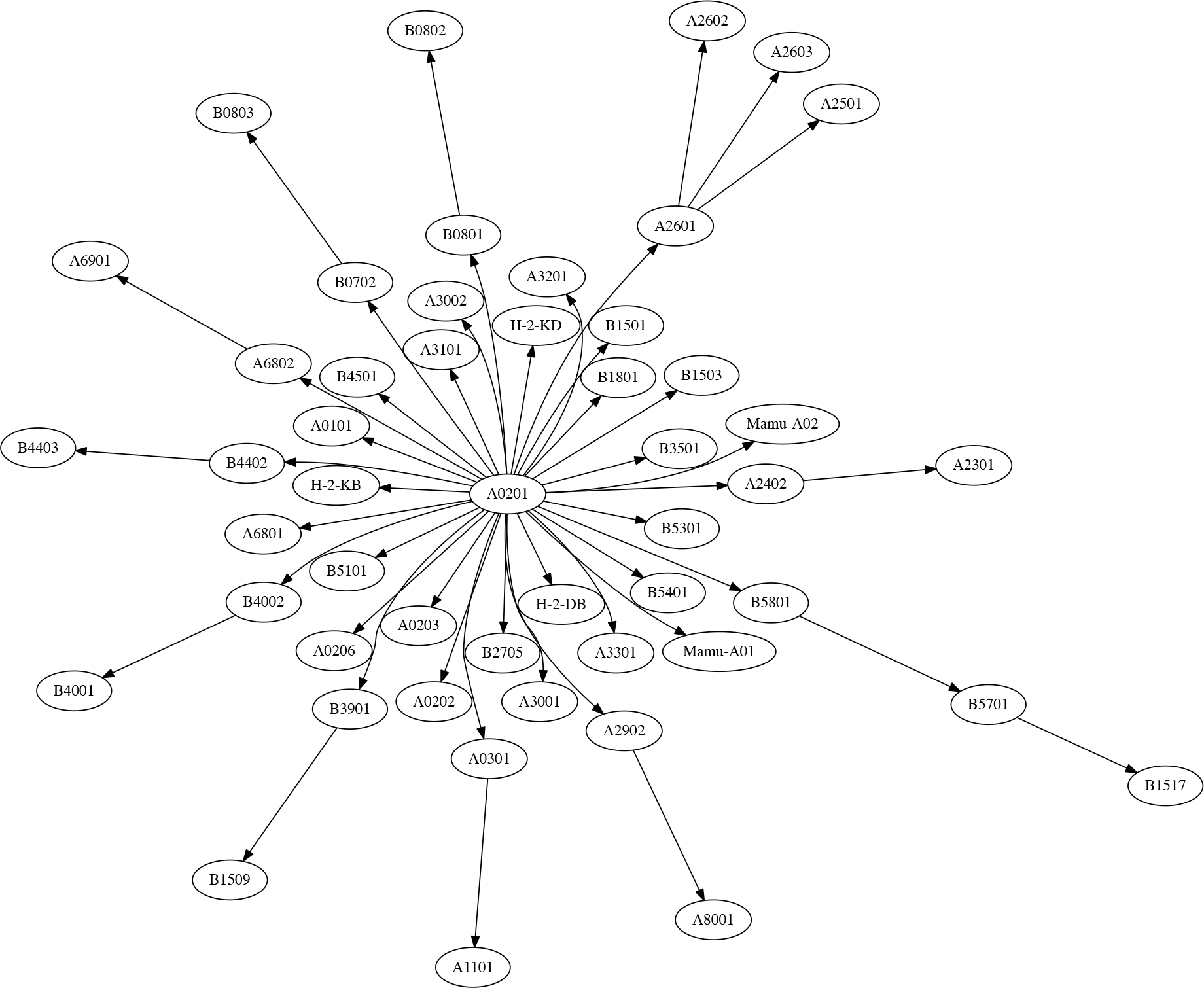
Transfer learning directed graph. The graph shows transfer of information between MHC Class I alleles in our network training protocol. Nodes represent MHC class I allele-specific neural networks and directed edges show how trained network weights are transfered from one network to another. The first network trained is for HLA-A*02:01 binding peptides, and the trained weights are used to initalize the networks for all other alleles, yielding 50 additional allele-specific networks. These networks are tested for their prediction performance on the training data for each of the 50 alleles. The network weights for the best performing networks are transferred to the corresponding alleles. This three-step protocol is shown in Figure S2.

**Figure S2.**
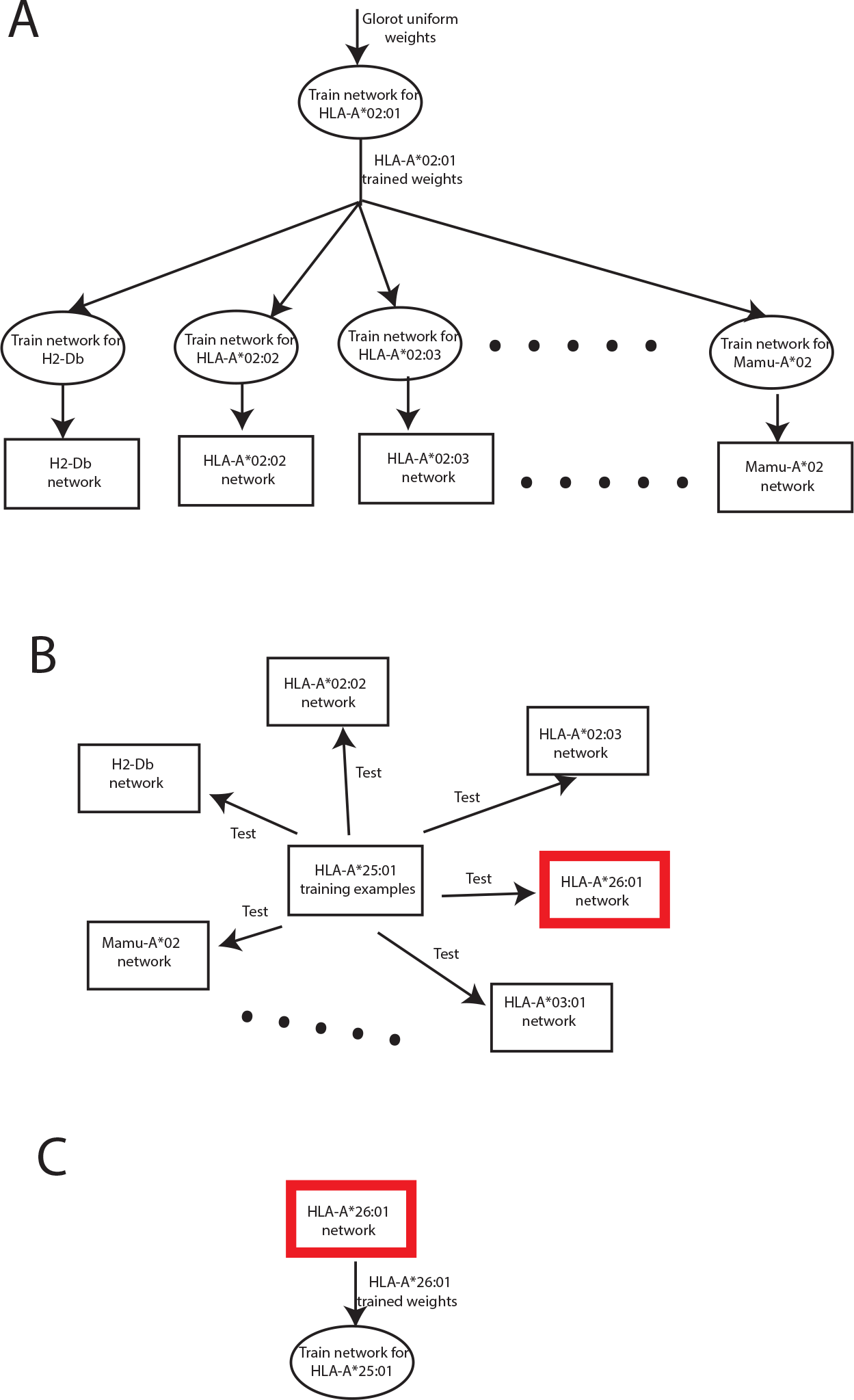
Three steps of the transfer learning protocol. **A.** Network weights of HLA-A*02:01 are used to initialize 50 networks for all other alleles in the benchmark. **B.** The training examples for each allele are tested for performance using all the networks trained in A. Example for a single allele HLA-A*25:01 is shown. The network with the highest AUC (originally trained on HLA-A*26:01) is selected (red highlight). **C.** The network weights for the best networks found in the previous step (e.g., HLA-A*26:01) are used to initialize and retrain the network for each allele of interest.

**Figure S3.**
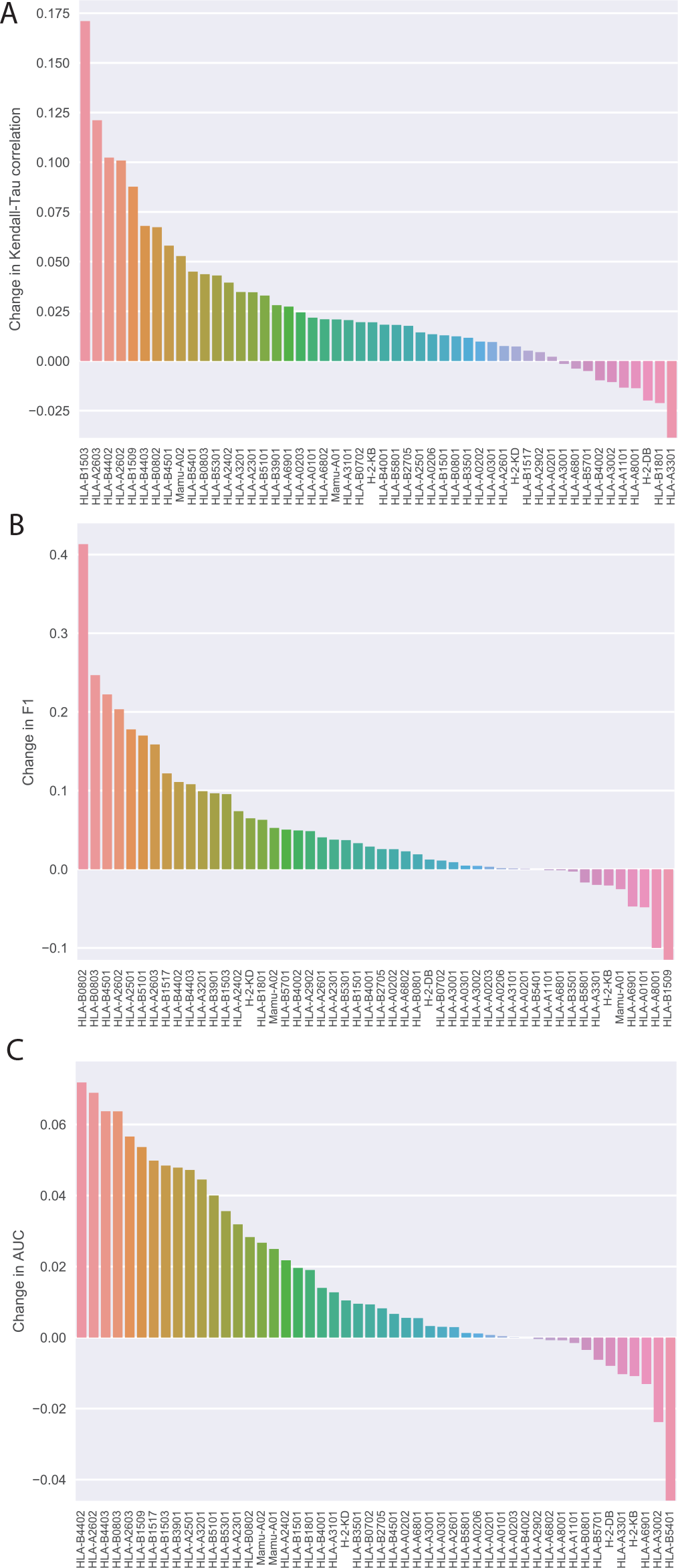
Transfer learning improves prediction peformance. **(A)** Kendall-Tau correlation. **(b)** F1. **(c)** AUC. Comparison is between MHCnuggets trained with and without the transfer learning protocol.

